# The allergic response mediated by fire ant venom proteins

**DOI:** 10.1101/382697

**Authors:** Zamith-Miranda Daniel, Eduardo G. P. Fox, Ana Paula Monteiro, Diogo Gama, Luiz E. Poublan, Almair Ferreira de Araujo, Maria F. C. Araujo, Georgia C. Atella, Ednildo A. Machado, Bruno L. Diaz

## Abstract

Fire ants are widely studied, invasive and venomous arthropod pests. There is significant biomedical interest in immunotherapy against fire ant stings. However, mainly due to practical reasons, the physiological effects of envenomation has remained poorly characterized. The present study takes advantage of a recently-described venom protein extract to delineate the immunological pathways underlying the allergic reaction to fire ant venom toxins. Mice were injected with controlled doses of venom protein extract. Following sensitization and a second exposure, a marked footpad swelling was observed. Based on eosinophil recruitment and production of Th2 cytokines, we hereby establish that fire ant proteins *per se* can lead to an allergic response, which casts a new light into the mechanism of action of these toxins.

## Introduction

The Red Imported Fire Ant (RIFA) *Solenopsis invicta* Buren (Insecta: Formicidae) is one of the most dangerous invasive pests on a global scale^1,2^. These aggressive ants have been inadvertently shipped from South America to many territories around the world over the last century, and are now causing severe problems in regions as far apart as Vietnam, China, Australia, the United States, and the Galapagos archipelago^1,3^. When disturbed, these ants will viciously defend their nests with a venomous sting which can be particularly dangerous to the children^4^ and the elderly^5^.

Anaphylactic reactions to fire ant stings are recurrent for a fraction of previously sensitized victims, where systemic hypersensitive reactions can pose life-threatening complications^6,7^. Since the prevalence of sensitized subjects in invaded areas is reportedly relatively high^8^ there is a growing demand for immunotherapy methods in regions recently invaded by RIFA^9,10^. Immediate effects of the stings are mainly caused by a major (> 95%) fraction of toxic alkaloids, but the later allergic responses^11,12^ are solely ascribed to venom proteins.

Nonetheless, in spite of over 40 years of research^13^, the allergic reaction caused by fire ant venom has remained poorly characterized. This is primarily because each ant carries venom proteins that amounts to only 0.1% of total body weight^14,15^, rendering bioassays unfeasible. Fortunately, a simple method enabling the isolation of venom proteins from fire ants in large quantities has been recently devised^14^, enabling a range of biological tests which were previously impossible, such as injecting animal models with venom protein fractions.

The network of immune cells and expressed factors involved in an allergic response is complex and context-dependent. We provide a simplified general diagram illustrating the proposed allergic reaction induced by injected venom in Figure 1. Allergens upon entering the system are endocytosed and processed by phagocytes (e.g. macrophages, dendritic cells) in order to generate peptides. The physicochemical properties and enzymatic activity of allergens play a central role in the activation and maturation of dendritic cells, which are fundamental steps for proper antigen presentation and immune response triggering. The dendritic cells subsequently migrate to the nearest draining lymph node to present the generated peptides (antigens) coupled to a major histocompatibility complex class II (MHCII) molecule to naive T lymphocytes^16^. Three signals are required during the antigen presentation in order to induce proper T cell activation: the first is provided by the recognition of the complex MHCII/peptide by the T cell receptor (TCR) present in a naive T lymphocyte. The binding between the phagocyte costimulatory molecules (e.g. CD86) and the CD28 receptor present at the T cell surface provides the second signal. The third signal is given by soluble cytokines released from the antigen presenting cell (APC) to act on the T cell. Together, these signals can activate a naive T cell that will differentiate into distinct T subsets^17,18^.

**Figure 1.**
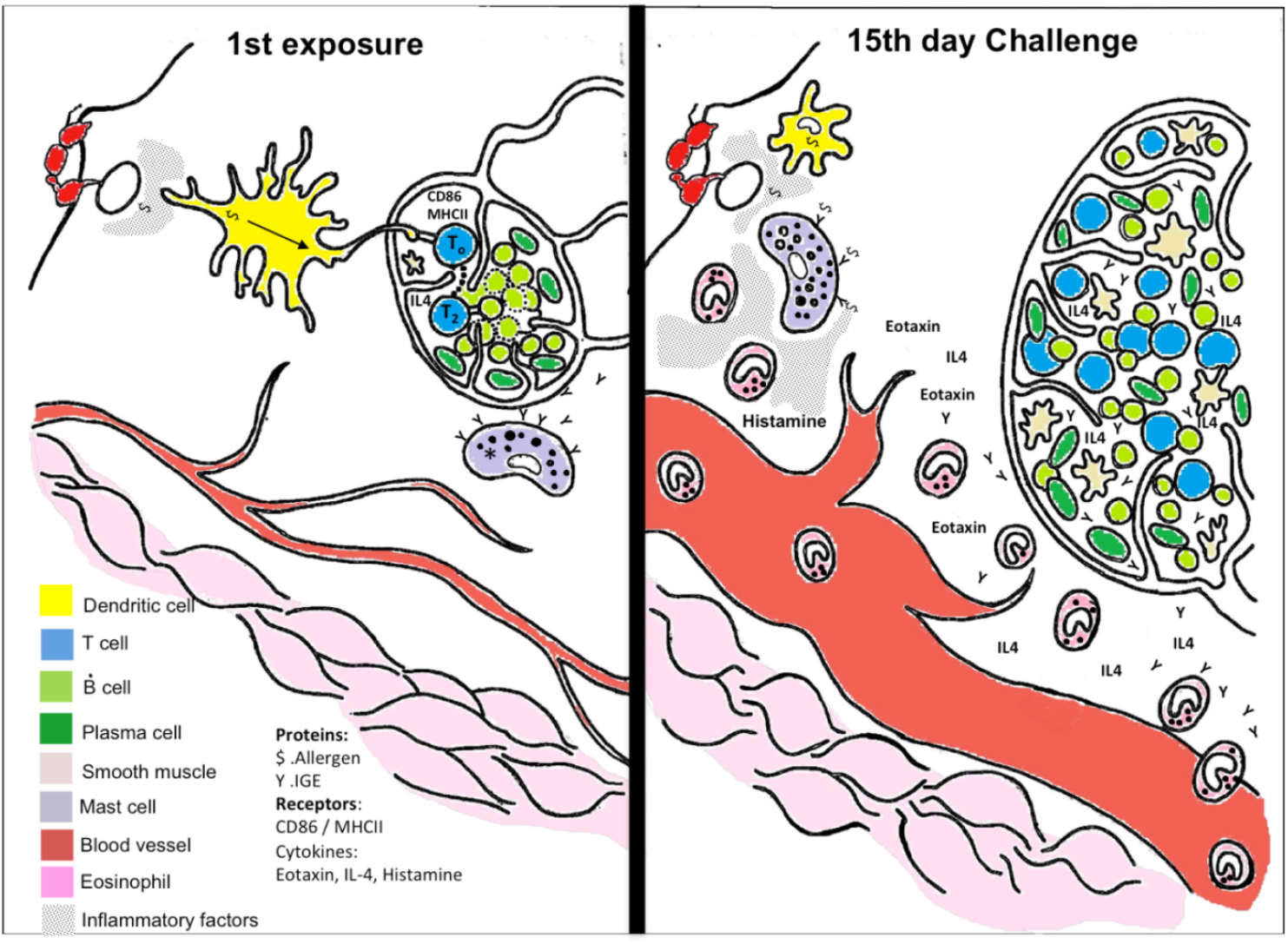
Hypothesized allergic reaction following a fire ant sting. Upon a first exposure, (left panel) peripheral resident cells recruit blood leukocytes that will differentiate into macrophages and dendritic cells. These phagocytes will endocytose and process the venom’s antigens. Dendritic cells exposed to venom components will be activated and mature, thus promoting their ability to trigger the adaptive immune response. In this scheme, a dendritic cell migrates into a lymph node to present fire ant venom-derived peptides via MHCII to a naive T cell in the presence of the co-stimulatory molecule CD86. Upon activation, T cells become TH2 cells and proliferate under secretion of IL-4, and which will eventually activate an antigen-compatible B lymphocyte. Activated lymphocytes undergo clonal expansion inside the lymph node. A key event at the immunization stage takes place within ca. 7–10 days: activated B cells mature into IgE-secretory plasma cells, and the secreted allergen-specific IgE bind to peripheral mast-cells’ FcɛR (this immunization event is marked with an *). Upon a second exposure (right panel), antigens will trigger a specific, amplified reaction: (i) mast cells carrying antigen-specific IgE are activated to secrete inflammatory factors (not tested in the present study); (ii)specific lymphocytes secrete IgE and IL-4 promoting further cell activation; (iii) the amplified reaction increases edema and promote intense recruiting of eosinophils.

In the context of allergy, the participation of type-2 T helper cells (Th2) is central. Once activated, Th2 lymphocytes will undergo clonal expansion leading to lymph node hyperplasia and production of IL-4, the major Th2 cytokine^19,20^. In the context of venoms, many toxins can directly trigger Th2 responses^21,22,23,24^. This T cell subset produces cytokines that can promote leukocyte recruitment and the synthesis of further Th2-related cytokines^17^. A second exposure to venom antigens triggers a stereotypical allergic response marked by acute vascular permeability and later eosinophil recruitment at the site of the sting. This condition can be life threatening when the allergic reaction is triggered systemically progressing to anaphylactic shock.^22^

In this context, the present manuscript utilizes an extract of RIFA venom proteins to describe the general features of the immunological response from injections into male mice. These venom proteins are demonstrated to be independent from alkaloids and adjuvants in triggering an immunological response, which brings a new biological meaning to the role of these toxins.

## Results and Discussion

In short, all challenged mice demonstrated clear signs of allergic response, demonstrating the ant venom proteins will immunize even at the lowest tested doses, and in the absence of adjuvants upon first exposure. Details are given as per the pertinent sections below.

## Ant venom promotes eosinophil recruitment to peritoneal cavity

To test the premise that venom proteins are causative factors of allergic reactions to fire ant stings, we sensitized mice via venom proteins into the hind footpad. 14 days later, the same mice were challenged by a second injection into the peritoneal cavity. The observed increase in peritoneal cellularity in venom-sensitized mice was mainly due to significant eosinophil recruitment. Injection of venom proteins into non-sensitized (naive) mice promotes merely mild, non-significant eosinophil recruitment (Fig. 2). Interestingly, inoculating ovalbumin (OVA) into mice previously sensitized with combined venom proteins + OVA promotes recruitment of eosinophils comparable to positive controls of combined OVA + aluminum hydroxide (as an adjuvant). This result suggests a potential role of venom proteins as adjuvants in allergic sensitization.

**Figure 2.**
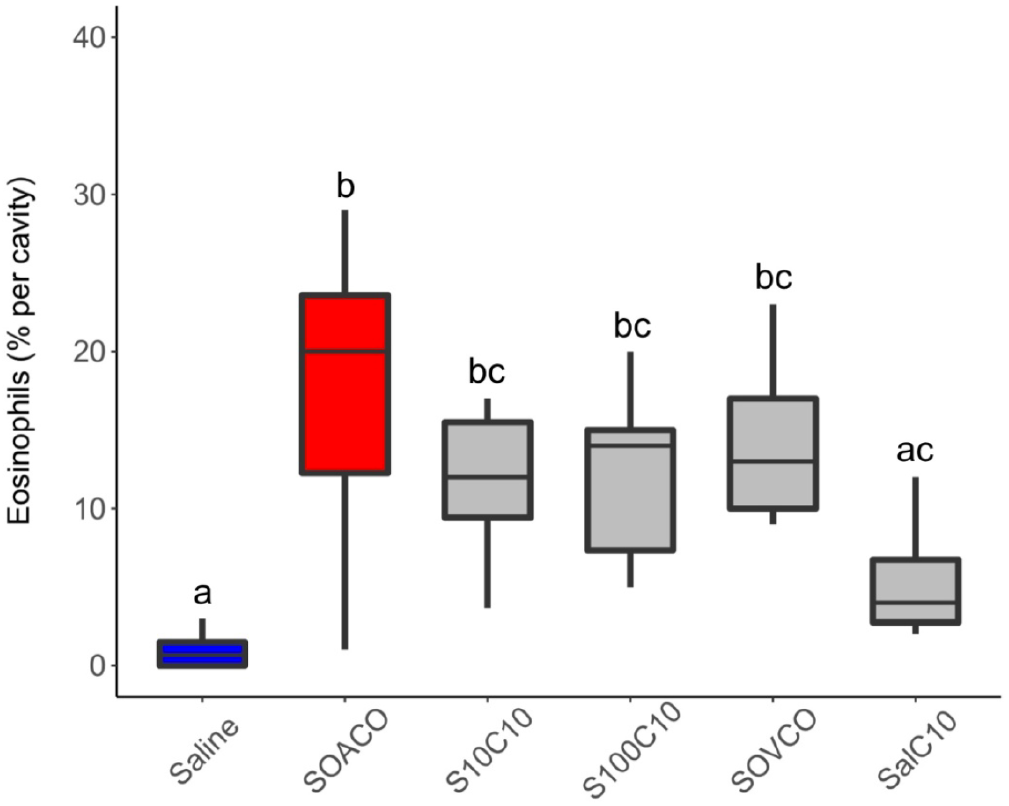
Ant venom promotes eosinophil recruitment to peritoneal cavity. Mice were previously
injected (sensitized: S) with saline (Sal), OVA + AlOH_3_ (SOACO), ant venom (10 μg = S10; 100 μg =S100), or OVA + 10 ¼g ant venom (SOVCO). After 2 weeks, mice were submitted to an intraperitoneal challenge (Challenge: C) with saline, OVA (CO), or 10 ¼g ant venom extract (C10). Twenty-four hours after the challenge, peritoneal cells were harvested and the eosinophils were counted. Measurements are expressed as the percentage of eosinophils among the peritoneal cell population. Boxplot of pooled internal replicates from three independent experiments, where the vertical lines are upper and lower limits, and the internal line are median values. Different letters indicate statistical significance (pvalue<0.05) between groups as compared by Kruskal-Wallis followed by Dunn’s Multiple Comparisons Test.

## Eotaxin production triggered by ant venom

Peritoneal eotaxin accumulation in mice challenged with venom protein was higher in saline-injected mice (Fig. 3). The increased eotaxin levels of venom inoculated mice were comparable to concomitant positive controls (see grey box ‘SOACO’ on Fig. 3). Taken together, Figures 2 and 3 strongly suggest that venom proteins *per se* can actively promote sensitization-dependent eosinophil recruitment to the peritoneal cavity after a second exposure through the production of eosinophilotactic signals such as eotaxin.

**Figure 3.**
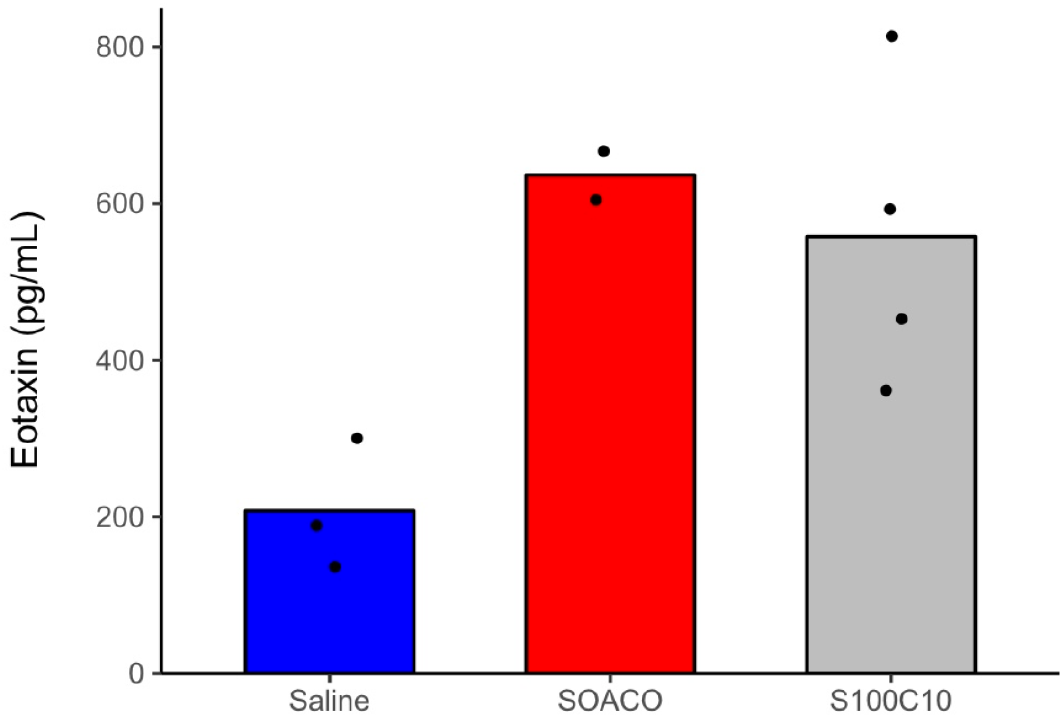
Fire ant venom induces eotaxin production in the peritoneal cavity. Mice were previously immunized with saline, ovalbumin + Al(OH)_3_ (SOACO), or 100 μg ant venom extract (S100C10). After 2 weeks, mice were submitted to an intraperitoneal challenge with saline, ovalbumin,or 10 μg ant venom extract, respectively. Within 24 h after the challenge, mice peritoneal exudates were harvested and eotaxin quantification was performed by ELISA. Axis numbers are calculated concentrations of eotaxin in pg/mL. Bars represent means from pooled replicates deriving from two independent experiments; dots are raw values. Treatment S100C10 was statistically different from Saline control by nonparametric Wilcoxon-Mann-Whitney test at alpha = 0.05.

## Dendritic cells activation by ant venom

To investigate a potential role of RIFA venom proteins as adjuvants in immunization, we evaluated murine bone marrow-derived dendritic cells (BMDC) after exposure to venom proteins or alkaloids from the venom (Fig. 4 A-C). Dendritic cells are responsible for the induction of adaptive immune responses, and seen as ‘sentinels’ for the immune system^25^. The venom alkaloids proved extremely cytotoxic even at 10 μg/mL (data not shown) indicating these venom components may not be relevant direct immunogenic stimuli for dendritic cells. Meanwhile, the venom proteins elicited an activation response, as shown by the upregulation of MHCII and CD86 (Fig. 4). This demonstrates that the venom proteins can directly activate dendritic cells to promote antigen presentation, of which the increased expression of MHCII is a first marker signal of immunological activation. Following expression of MHCII, secondary signals (costimulatory signals) are needed to elicit a response from lymphocytes, including CD86. The fact that the highest doses of venom generated a weaker response may indicate the presence of intrinsically dynamic components, such as proteinase-regulated zymogens and enzymatic inhibitors. Toxin activation by dilution could be a simple mechanism to achieve high levels of damage towards injected victims whilst ensuring natural resistance by the ants. In fact at least one phospholipase inhibitor is described from the venom of RIFA^26^, indicating secreted proteases might be involved in the activation of zymogens.

**Figure 4.**
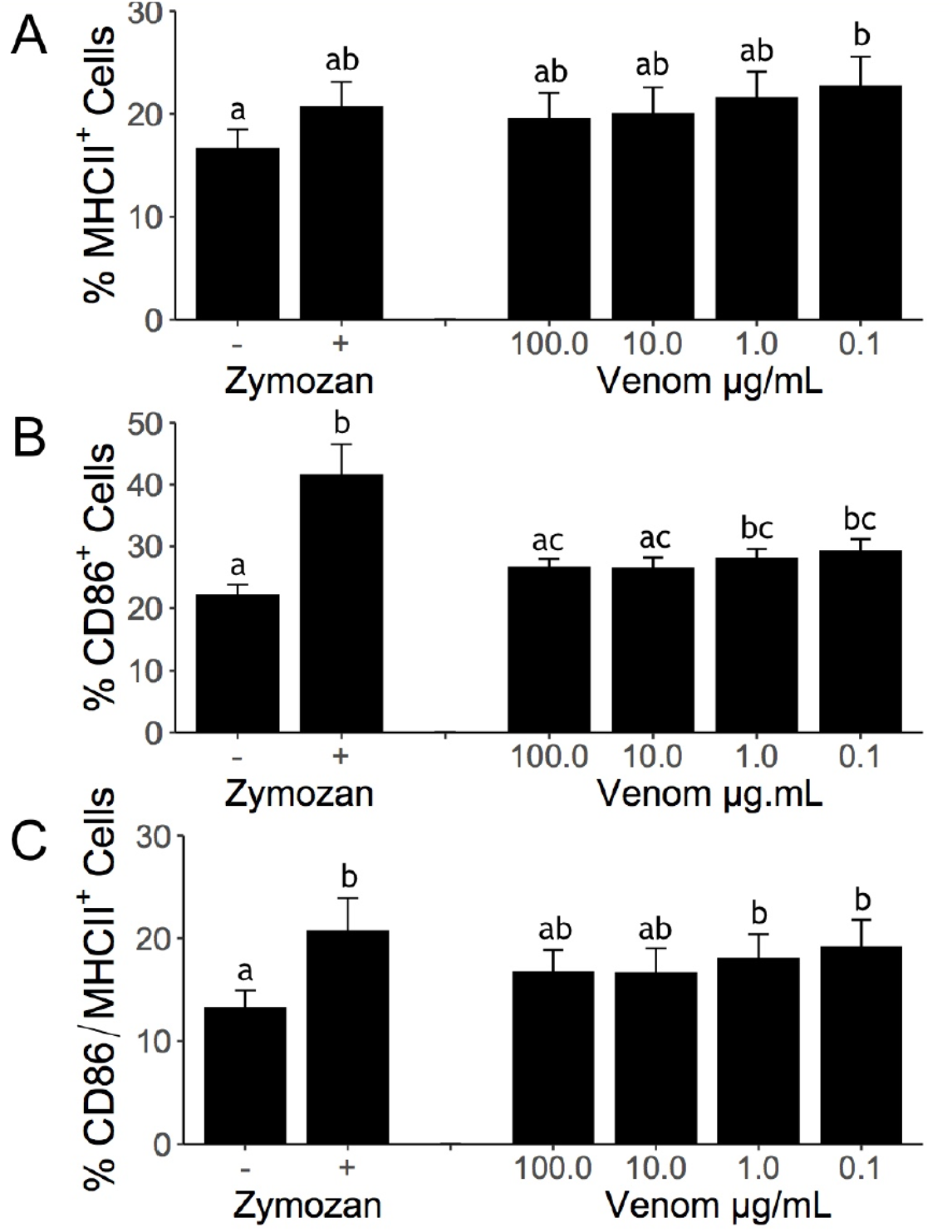
Ant venom activates dendritic cells in vitro. BMDCs were stimulated with zymosan (1:5)or with different concentrations of fire ant venom extract. After a 24h incubation period, the expression of MHCII (A), CD86 (B), and MHCII+CD86 (C) were evaluated. Bars represent means and errors of three independent experiments. Different letters indicate statistical significance (p-value<0.05) between groups as compared by Kruskal-Wallis followed by Dunn’s Multiple Comparisons Test.

## *In vivo* hypersensitivity

The literature reports that the hypersensitive reactions following RIFA stings may include swelling of distal body parts which might lead to a life-threatening throat obstruction in humans^6,7^, initially signaled by wheezing, difficulty breathing, and a coarse voice. To test for allergen-triggered swelling, mice were sensitized with an intradermal inoculum of venom proteins into their right hind footpad (sensitization), and 14 days later the opposite hind footpad (left) was challenged with another inoculum. Both footpads were measured every 30 min from the moment of inoculation until up to 2h after injections.

Naturally, all assayed mice displayed marked swelling in the injected footpad, as a result of the inoculum, but the effects become visible as the injected volume is drained. Full drainage of the initial swelling is completed within 2 hours following the injections. After 2 weeks following the first exposure, a second inoculation (i.e. challenge) resulted in persistent footpad swelling of venom-exposed animals comparable to positive controls (OVA + Al_2_(OH)_3_) (Fig. 5). To address a possible role for venom proteins as adjuvants, mice were sensitized with an association between OVA and venom protein, or just OVA. When further challenged with OVA, only the mice that received OVA + venom proteins presented footpad swelling (Fig. 6). These results establish a sensitization-dependent swelling upon later challenge as an immunogenic response to fire ant venom proteins.

**Figure 5.**
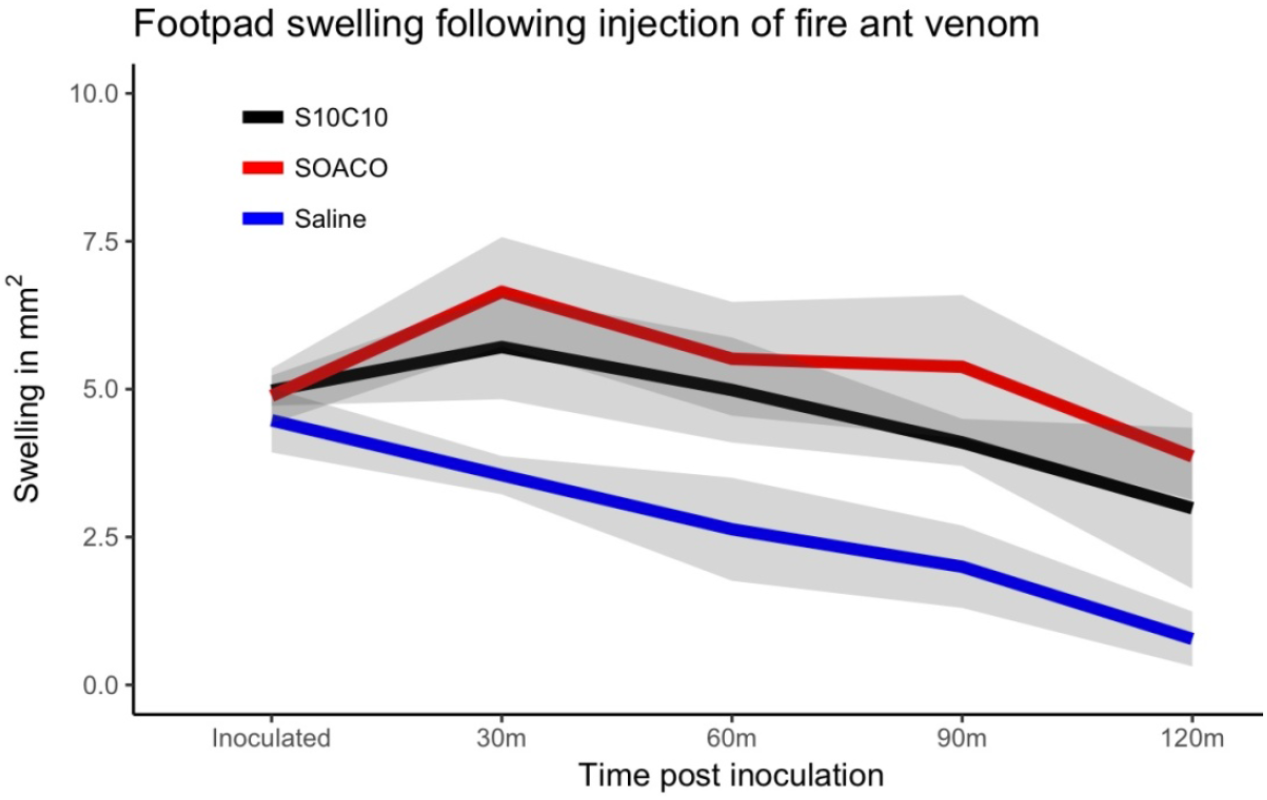
Footpad swelling after ant venom protein-fraction challenge. Mice were previously
sensitized with saline, ovalbumin (OVA) + AlOH_3_ (SOACO), or fire ant venom extract (S10C10). After 2 weeks mice were respectively challenged with another exposure to saline, OVA or ant venom extract, and their footpad swelling was measured for 120 minutes. Lines represent means of the obtained swelling measures and the shaded area are standard errors from three independent experiments (N = 5 mice per group from 3 independent experiments).

**Figure 6.**
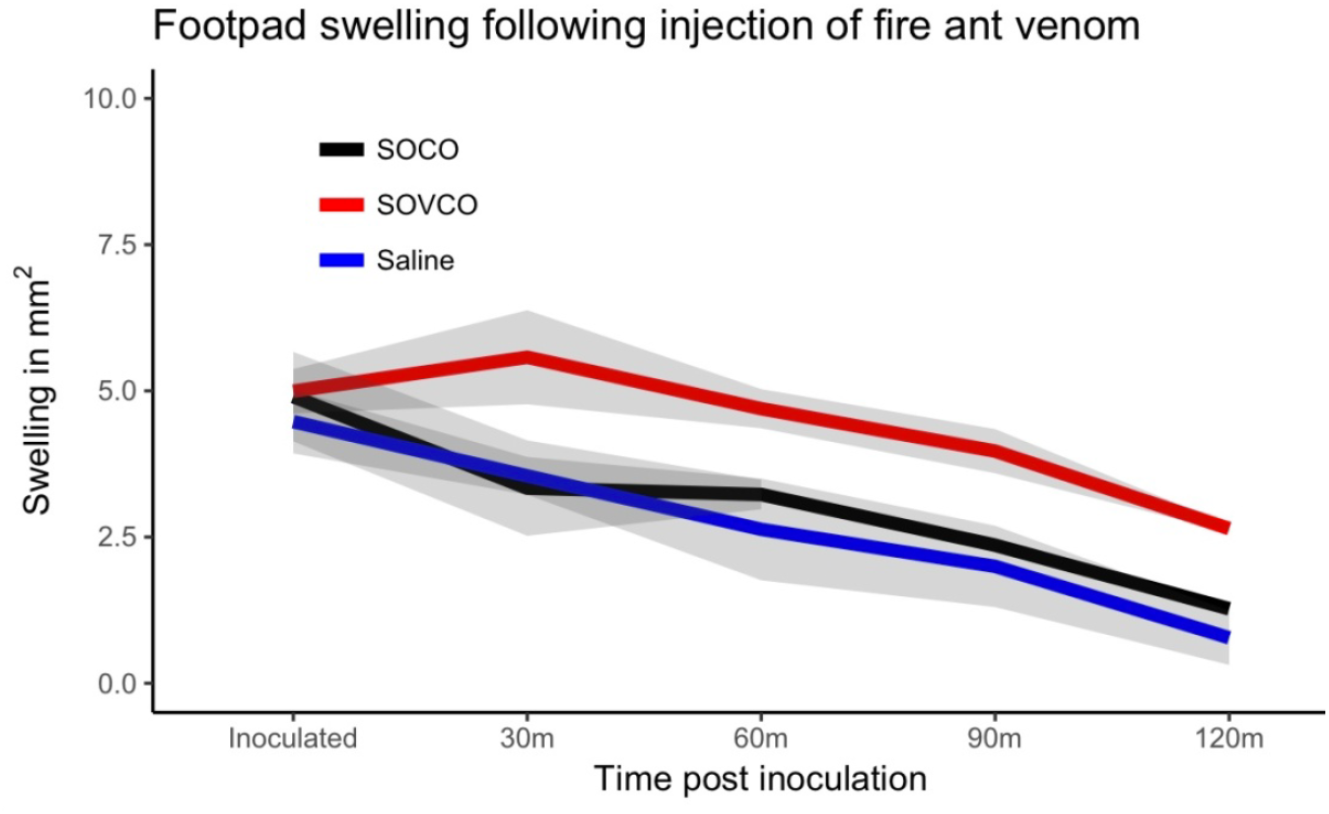
Adjuvant function of fire ant venom proteins. Mice were previously immunized with saline, ovalbumin + ant venom extract (SOVCO), or ovalbumin (SOCO). After 2 weeks the mice were challenged with new injections of saline or ovalbumin; footpad swelling was measured within 30–120 minutes. Lines represent means; shaded area are standard errors measured at each analysis time point (N = 5 mice per group from 3 independent experiments).

## Draining lymph node response

We assessed the hyperplasia of the draining lymph node (popliteal) 14 days after sensitization (Fig. 7). The lymph node of positive control mice (OVA + Al(OH)_3_) became greatly swollen two weeks after the sensitization. Venom-sensitized mice showed marked but less intense lymph node hyperplasia when sensitized with venom proteins. These lymph nodes were dissected out of the euthanized mice to evaluate the number of cells and the production of IL-4, the major Th2 cytokine, in order to test whether venom proteins induced priming of T cells *in vivo.* As indicated by the size of the respective lymph nodes, venom protein-sensitized mice presented an increase in the total cell number per lymph node when compared to the negative controls. Interestingly, lymph node hyperplasia from mice sensitized with the highest dose of venom proteins (100 μg) was not statistically different from the negative control, although a clear trend is apparent (Fig. 7). The increased cell number is indicative of lymphocyte proliferation triggered by venom proteins exposure, although this proliferative activity was less intense than the positive control (OVA + Al(OH)_3_). However, when stimulated *in vitro* for cytokine production, cells from the draining lymph nodes of venom-sensitized mice produced significant amounts of IL-4 (Fig. 8), comparable to OVA + Al(OH)_3_ sensitized positive controls, with an apparent tendency to greater production after 48h (see whiskers on right hand side of axis on Fig. 8).

**Figure 7.**
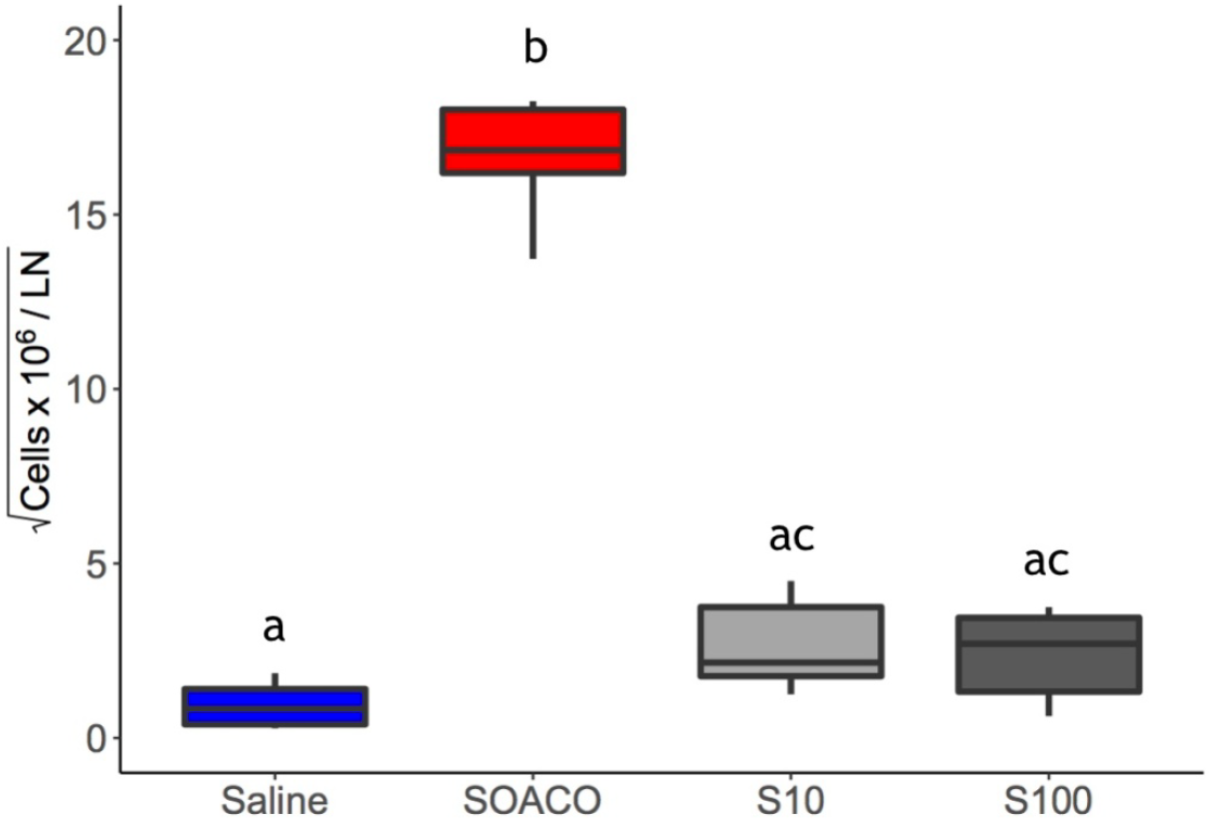
Fire ant venom induces lymph node response. Mice were previously immunized with either OVA + AlOH_3_ (SOA) or with 10 μg (S10) or 100 μg (S100) of of ant venom proteins extract. 2 weeks after first exposure, the draining (popliteal) lymph nodes (LN) were isolated and cell totals were determined. The box plot represents upper and lower limits and quartiles from six independent experimental results; the internal line is the median. Different letters indicate statistical significance (p < 0.05) between groups as compared by Kruskal-Wallis followed by Dunn’s Multiple Comparisons Test.

**Figure 8.**
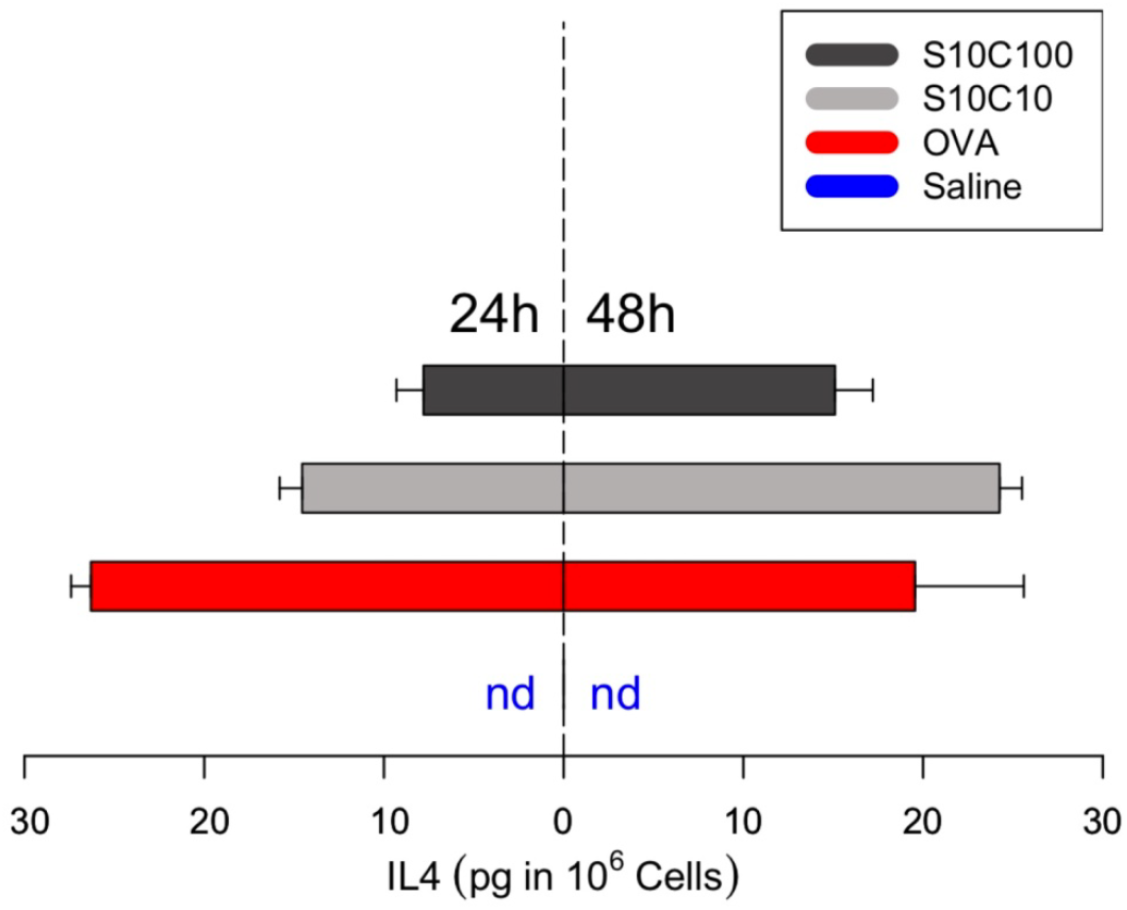
Ant venom induces cytokine response in lymph node cells. Mice were previously immunized with saline, ovalbumin + AlOH_3_ (SOA) or with 10 μg (S10) or 100 μg (S100) of ant venom proteins. After 2 weeks, popliteal lymph nodes were dissected and macerated for cell extraction, followed by stimulation with α-CD3. The cytokine IL-4 was quantified from lymph nodes supernatants after 24 h (left side) and 48 h (righthand side) of incubation. Bars are mirrored for size comparison representing the means(n = 2 independent experiments) while the whiskers are the interval towards the maximum value; ‘nd’ = not detectable.

<R scripts behind statistical analyses and plots available as supplementary data>

## Venom proteins as adjuvant

At least four different allergens are described from RIFA venom proteins^13^. However, the venom components driving the adaptive immune response have not been empirically identified. Crude fire ant venom is mainly composed of alkaloids, which are cytotoxic and insoluble^13,26^ and cause local tissue damage. Tissue damage is accompanied by a steep inflammatory response, typically followed by later pustule formation. Given the local inflammatory reaction and edema that follows fire ants stings, it is tempting to hypothesize that alkaloids would play a necessary biological role as adjuvants for ensuring the adaptive response (as in Fig. 1) to the few accompanying venom antigens, roughly akin to the role of aluminum adjuvants in injected vaccines^27,28^. However, venom alkaloids herein proved to be highly toxic to dendritic cells, while the proteins were sufficient to promote antigen presentation (increased MHC-II) and co-stimulatory (increased CD86) capacity (Fig. 4). To test this hypothesis, we injected naive mice with a mixture of venom proteins and ovalbumin in the absence of an aluminum adjuvant. Ovalbumin is innocuous when injected alone, but becomes a potent allergenic when administered in combination with an adjuvant like aluminum hydroxide (Fig. 5), thus widely employed as a control in immunology experiments.

Mice previously sensitized with 100 μg of ovalbumin in the presence of 10 μg of fire ant venom protein extract (**SOVCO**) produced a significant increase in footpad volume when later challenged with ovalbumin, as compared to challenged mice previously sensitized with ovalbumin alone (SOCO) (Fig. 6). The amount of swelling was similar to OVA + Al(OH)_3_ sensitized mice (SOACO) (Fig. 5). A similar adjuvant effect was observed when the OVA challenge injection was performed into the peritoneal cavity, where eosinophil influx was determined 24 h later (SOCO: 5.24 ± 1.53%; SOVCO: 12.15 ± 4.55%, p<0.05). Since the extraction of venom proteins is not specifically demonstrated to eliminate potential microbial contaminants (e.g. LPS) such compounds could be involved in the observed effects.

To assess this possibility, we designed additional assays with heat-treated venom. Injection of venom induced significant eosinophil recruitment only when animals were previously exposed to the venom, indicating that neither contaminants nor the venom extract itself will trigger eosinophilic response without a previous sensitization (Fig. 2). Still, a possible role of contaminants in sensitization remained. To address this point, mice were sensitized with 100.0 μg of OVA combined with 10.0 μg of previously heat inactivated (100 °C / 60 min) venom extract proteins (SOVhiCO). When SOVhiCO mice were challenged with OVA they elicited comparable amounts of eosinophils in the peritoneal cavity as mice sensitized with OVA alone (SOCO) (Fig. S2). Thus, heat exposure completely eliminated the adjuvant property of the venom extract, suggesting that it is fundamentally enzymatic, given that bacterial contaminants such as LPS are thermo-resistant^29^.

Fire ant venom proteins are relatively few in comparison to the venom composition of other social insects^26^. Still, some of the identified proteins may contribute to the observed adjuvant effects. Toxins identified from the venom extract include proteins involved in tissue damage (phospholipases A_1_ and A_2_, disintegrin/ metalloproteinase, and myotoxins), and neurotoxicity (toxin PsTX-60, U5-ctenotoxins Pk1a, alpha-toxins Tc48a, and *Scolopendra* toxin)^26^. Phospholipase A_2_ from honeybee venom is known as a potent inducer of activation and maturation of dendritic cells^30^, suggesting that this enzyme may be playing a similar role in RIFA venom. In fact, previous immunological tests with honeybee venom indicated that venom-purified phospholipase A_2_ alone could lead to an adaptive response via Th2 phenotype^31,32^. Activity of PLA_2_ activity in RIFA venom was herein demonstrated for the first time with a fluorogenic substrate assay^33^. Remarkably the assay performed with 10.0 μg of the venom protein yielded the same activity intensity as 13.0 μg of *Bothrops jararaca* venom (Fig. S1). These results also demonstrate that enzymatic activity of the venom is maintained through extraction, lyophilization, and storage.

## Conclusions

The present observations delineate the general pathways by which RIFA venom proteins can drive the primary physiological effects of an allergic response, i.e. the basic mechanisms of immunosensitization induced by fire ant venom proteins depicted in Fig. 1. Immunosensitization was manifested in all venom-exposed individuals, visually realized by footpad swelling. Peritoneal cell counts indicated a prototypical Th2 immune reaction based on augmented recruitment of eosinophils probably due to the production of eotaxin. The production of IL-4 by draining lymph node cells illustrates the cellular immune aspects of the allergic reaction. Co-injection of fire ant venom proteins with ovalbumin induced sensitization to ovalbumin, demonstrating an adjuvant activity for the venom itself decoupled of a pro-inflammatory effect of venom alkaloids.

From our standpoint such first insights into the immunoreaction to fire ant venom proteins, as tested in mice, pave the way for further tests. For example, it would be interesting to investigate individual and species-specific variability, and the response to purified venom fractions. Identifying the most immunologically active peptides is paramount for developing more cost-effective, safer immunotherapy strategies. The venom extraction technology^13,14^ could include further purification steps (e.g. gel exclusion chromatography) to test finer fractions in animal models towards validation for immunotherapy. As a next step, trying to revert the acquired sensitivity of venom-injected mice could provide a simple model for powerful studies of immunotherapeutic protocols. Moreover, other aspects involved in immune/allergic responses for RIFA (e.g. venom fractions) and others related species are under investigation.

## Methods

### Venom sample

Nests of RIFA were excavated from the public gardens at Ilha do Fundão, Rio de Janeiro, Brazil into a plastic bucket rimmed with Teflon paint, and the ants were separated from the soil by slow flooding^34^. Species identification was based on a distinctive medial clypeal tooth and clearly-defined frons mark present on major workers, apart from typical color pattern of workers and the distinctively black males^35^. Based on recurrent field observations of excavated nests in this region by EGPF, local RIFA populations seem to be consistently monogyne^3^ (i.e. low density of mounds, presence of single reproductive queen, and relatively large major workers). The ants were allowed to dry and cleanse for 2 hours, and then submitted to a rapid method of venom extraction^13–14^, which relies on immersing live ants in a biphasic solvent mix (distilled water : hexane at 1 : 5). The aqueous phase was subsequently lyophilized (henceforth ‘RIFA venom proteins’) and maintained at -20°C until reconstitution for use. RIFA venom proteins extract obtained through method^14^ is equivalent to commercially available preparations as demonstrated previously by 2D electrophoresis^14^ and mass spectrometry^26^. Upon each day of experimentation, a dedicated sample of RIFA venom proteins was reconstituted in saline (sterile 0.9 % NaCl) for fresh use.

### Venom injections and footpad swelling

Male *Mus musculus* BALB/c mice aged 8–12 weeks were reared at 25 °C and 70 % humidity in metal cages, and given water and special chow *ad libitum*. All methods used in this study were approved by the Ethics Committee on the Use of Animals of Health Sciences Center of Federal University of Rio de Janeiro, Brazil (CEUA/CCS/ UFRJ, CONCEA). All animals received humane care in compliance with the “Principles of Laboratory Animal Care” formulated by the National Society for Medical Research and the “Guide for the Care and Use of Laboratory Animals” prepared by the National Academy of Sciences, USA, and National Council for Controlling Animal Experimentation, Ministry of Science, Technology and Innovation (CONCEA/MCTI), Brazil. Males were always siblings (to avoid fights) being held at the number of four or five per cage. Hypodermic plastic syringes were used; all intradermal injections were performed on mice anesthetized with ketamine 112.5 mg/kg and xylazine 10.0 mg/kg unless stated otherwise. The first venom presentation (i.e. the sensitizing inoculum) was injected into the right hind footpad using either 10 or 100.0 *μ*g of total venom protein per animal. A third sensitization test was made using 10.0 μg of venom protein previously incubated at 100 °C in a water bath for 60 min, to test for enzymatic adjuvant role (SOVhiCO). Fourteen days after sensitization, a second exposure to venom (i.e. challenge) was made into the left hind footpad. This interval was established to allow for immunization of tested animals against the fire ant venom allergens, as detailed in Fig. 1 (see the asterisk on left-hand panel). Immediately before and after each venom injection, the width and height of the challenged and opposing paws were measured with a caliper, and measurements were taken every 30 min for 2h, during which animals were monitored. Negative control injections used saline solution (sensitization - S; and challenge - C), and positive controls (SOACO) used ovalbumin (OVA - 100.0 μg/animal) with adjuvant aluminum hydroxide (5.0 mg/animal) upon sensitization, or only OVA upon challenge. All footpad inoculations were carried out in a 30.0 μL injection. Experimental groups were sensitized and challenged with venom protein unless stated otherwise. Each experiment employed groups of 4–5 mice and were repeated at least 3 times independently, unless specified otherwise.

### Leukocyte recruitment to the peritoneal cavity

Sensitized mice were intraperitoneally challenged with 0.9% sterile saline, OVA, or venom proteins with the quantities of OVA and venom previously described for a total injection volume of 500.0 μL per individual. Twenty-four hours after the challenge, peritoneal cavities were washed with PBS containing 0.1% heparin and the resulting suspensions were used for total and differential cell counting, using a Neubauer chamber and cytospun slides, respectively. The slides were stained with panoptic stain. The peritoneal exudates were harvested for chemokine quantification (Eotaxin - BDBiosciences) by ELISA according to the manufacturer’s protocol.

### Lymph node cytokine production

Mice were individually euthanized in CO2 chambers and the draining lymph node (popliteal) of the right hind footpad was dissected out and homogenized. Lymph node cells were cultivated for 48h in the presence of immobilized α-CD3 (1.0 μg/mL) for cytokine production. After the incubation period, supernatants were harvested and used to quantify IL-4 (BD Biosciences) by ELISA according to the manufacturer’s protocols.

### Phospholipase A2 (PLA2) activity in venom

PLA_2_ activity was measured using the fluorogenic substrate 1-acyl-2-{12-[(7-nitro-2-1,3-benzoxadiazol-4-yl) amino] dodecanoyl}-sn-glycero-3-phosphocholine (aka ‘12NBD PC’, Avanti Polar Lipids Inc), an analogous of phosphatidilcholine. Assays were done in 96 well plates in a medium composed of Tris-HCl buffer 20 mM, pH 7.5, CaCh 2.0 mM and 10.0 μg of venom protein of *S. invicta.* The reaction was triggered by the addition of 20.0 μM of the substrate 12NBD PC; after which the release of fatty acids was measured for 1 h in a plate-reading fluorimeter (Victor™ X5 Multilabel-Perkin Elmer) set for excitation and emission wavelengths of 460 nm and 534 nm, respectively. The negative control was reaction medium without added venom, and the positive control sample contained 1.0 μl (007E;13.0 μg/μl) of *Bothrops jararaca* venom^36^.

### Statistical Analyses

All statistical analyses and plots were generated with RStudio v.1.0.136, using the packages ‘plyr’, ‘reshape2’, ‘ggplot2’, ‘dunn.test’. All numeric raw data are provided as Supplementary Materials along with the scripts, to ensure output reproducibility and verification of specific details. Results from different internal and independent replicates were either compared for statistical differences using non-parametric Kruskal-Wallis followed by multiple comparison Dunn’s test (in comparing multiple treatments) or by Wilcoxon-Mann-Whitney test (in comparing two treatments). Our conclusions were also compatible with analytical patterns obtained by parametric methods (i.e. assuming Gaussian distribution; not shown).

## Acknowledgments

We wish to thank Leticia Coli and Debora Fogel for providing access to liquid nitrogen and lyophilizers; to the responsible technicians who provided the animals used in this manuscript. Eric Lucas has kindly revised a final version of this manuscript for readability and provided useful suggestions. Venom extract from *B. jararaca* was provided by André Lopes Fuly.

## Authors contributions

D.Z.M., E.G.P.F., E.A.M. and B.L.D. designed research; B.L.D. supervised the study;D.Z.M.,E.G.P.F., A.P.M, A.F.A., D.G and L.E.P. performed experiments; D.Z.M.,E.G.P.F.,A.P.M, D.G, L.E.P., E.A.M. and B.L.D. analyzed data; E.A.M. and B.L.D. contributed with new reagents/analytic tools; D.Z.M. and E.G.P.F. wrote the paper; E.A.M. and B.L.D. provided lab space, instrumentation and funding; all authors edited and gave critical input on the manuscript.

## References

1 Ascunce, M. S., Yang, C. C. J., Oakey, Calcaterra, L., Wu, W. J., Shih, C. J., Goudet, J., Ross, K. G., & Shoemaker, D. 2011. Global invasion history of the fire ant *Solenopsis invicta*. Science 331, 1066–1068, doi:10.1126/science. 1198734 (2011).

2 Morrison, L. W., Porter, S. D., Daniels, E. & Korzukhin, M. D. Potential global range expansion of the invasive fire ant, *Solenopsis invicta*. Biological Invasions 6, 183–191, doi:10.1023/b:binv.0000022135.96042.90 (2004).

3 Tschinkel, W. R. The fire ants. (Harvard University Press, 2006).

4 More, D. R., Kohlmeier, R. E. & Hoffman, D. R. Fatal anaphylaxis to indoor native fire ant stings in an infant. The American journal of forensic medicine and pathology 29, 62–63, doi:10.1097/PAF.0b013e3181651b53 (2008).

5 de Shazo, R. D., Williams, D. F. & Moak, E. S. Fire ant attacks on residents in health care facilities: A report of two cases. Annals of Internal Medicine 131, 424–429, doi:10.7326/0003-4819-131-6-199909210-00005 (1999).

6 Stafford, C. T. Hypersensitivity to Fire Ant Venom. Annals of Allergy, Asthma & Immunology 77, 87–99, doi:http://dx.doi.org/10.1016/S1081-1206(10)63493-X (1996).

7 Prahlow, J. A. & Barnard, J. J. Fatal Anaphylaxis Due to Fire Ant Stings. The American journal of forensic medicine and pathology 19, 137–142 (1998).

8 Partridge, M. E., Blackwood, W., Hamilton, R. G., Ford, J., Young, P., & Ownby, D. R. Prevalence of allergic sensitization to imported fire ants in children living in an endemic region of the southeastern United States. Annals of Allergy, Asthma & Immunology 100, 54–58, doi:http://dx.doi.org/10.1016/S1081-1206(10)60405-X (2008).

9 Potiwat, R., Tanyaratsrisakul, S., Maneewatchararangsri, S., Manuyakorn, W., Rerkpattanapipat, T., Samung, Y., Sirivichayakul, C., Chaicumpa, W., & Sitcharungsi, R. *Solenopsis geminata* (tropical fire ant) anaphylaxis among Thai patients: its allergens and specific IgE-reactivity. Asian Pac J Allergy Immunol. Online Earlier. doi:10.12932/AP-100217-0012

10 Haymore, B. R., McCoy, R. L. & Nelson, M. R. Imported fire ant immunotherapy prescribing patterns in a large health care system during a 17- year period. Annals of Allergy, Asthma & Immunology 102, 422–425, doi:http://dx.doi.org/10.1016/S1081-1206(10)60515-7 (2009).

11 Fitzgerald, K. T. & Flood, A. A. Hymenoptera stings. Clinical techniques in small animal practice 21, 194–204, doi:10.1053/j.ctsap.2006.10.002 (2006).

12 Parrino, J., Kandawalla, N. M. & Lockey, R. F. Treatment of local skin response to imported fire ant sting. Southern medical journal 74, 1361–1364 (1981).

13 Fox, E. G. P. in Venom Genomics and Proteomics (eds P. Gopalakrishnakone & Juan J. Calvete) 149–167 (Springer Netherlands, 2016).

14 Fox, E. G. P., Solis, D. R., dos Santos, L. D., dos Santos Pinto, J. R. A., da Silva Menegasso, A. R., Silva, R. C. M. C., Palma, M.S., Bueno, O. C., & Alcântara-Machado, E. A simple, rapid method for the extraction of whole fire ant venom (Insecta: Formicidae: *Solenopsis*). Toxicon 65, 5–8, doi:10.1016/j.toxicon.2012.12.009 (2013).

15 Baer, H., Liu, T.Y., Anderson, M.C., Blum, M., Schmid, W. H., & James, F. J. Protein components of fire ant venom (*Solenopsis invicta*). Toxicon 17, 397–405, doi:http://dx.doi.org/10.1016/0041-0101(79)90267-8 (1979).

16 Fujii, S., Liu, K., Smith, C., Bonito, A. J. & Steinman, R. M. The linkage of innate to adaptive immunity via maturing dendritic cells in vivo requires CD40 ligation in addition to antigen presentation and CD80/86 costimulation. The Journal of experimental medicine 199, 1607–1618, doi:10.1084/jem.20040317 (2004).

17 Zhu, J. & Paul, W. E. CD4 T cells: fates, functions, and faults. Blood 112, 1557–1569, doi:10.1182/blood-2008-05-078154 (2008).

18 Murphy, K. M. & Stockinger, B. Effector T cell plasticity: flexibility in the face of changing circumstances. Nature immunology 11, 674–680, doi:10.1038/ni.1899 (2010).

19 Kaplan, M. H., Schindler, U., Smiley, S. T. & Grusby, M. J. Stat6 is required for mediating responses to IL-4 and for development of Th2 cells. Immunity 4, 313–319 (1996).

20 Zheng, W. & Flavell, R. A. The transcription factor GATA-3 is necessary and sufficient for Th2 cytokine gene expression in CD4 T cells. Cell 89, 587–596 (1997).

21 Habermann, E. Bee and Wasp Venoms. Science 177, 314–322 (1972).

22 Bircher, A. J. Systemic immediate allergic reactions to arthropod stings and bites. Dermatology 210, 119–127, doi:10.1159/000082567 (2005).

23 Madero, M. F., Gámez, C., Madero, M.A., Fernández-Nieto, M., Sastre, J., & del Pozo, V. Characterization of allergens in four South American snake species. International archives of allergy and immunology 150, 307–310, doi:10.1159/000222684 (2009).

24 Muller, U. R. Insect venoms. Chemical immunology and allergy 95, 141–156, doi:10.1159/000315948 (2010).

25 Kapsenberg, M. L. Dendritic-cell control of pathogen-driven T-cell polarization. Nature reviews. Immunology 3, 984–993, doi:10.1038/nri1246 (2003).

26 Pinto, J. R. A. S., Fox, E. G. P., Saidemberg, D. M., Santos, L. D., da Silva Menegasso, A. R., Costa-Manso, E., Machado, E. A., Bueno, O. C., & Palma, M. S. Proteomic view of the venom from the fire ant *Solenopsis invicta* Buren. Journal of proteome research 11, 4643–4653, doi:10.1021/pr300451g (2012).

27 Hogenesch, H. Mechanism of immunopotentiation and safety of aluminum adjuvants. Frontiers in immunology 3, 406, doi:10.3389/fimmu.2012.00406 (2012).

28 Ghimire, T. R. The mechanisms of action of vaccines containing aluminum adjuvants: an in vitro vs in vivo paradigm. SpringerPlus 4, 181, doi:10.1186/s40064-015-0972-0 (2015).

29 Gorbet, M. B., & Sefton, M. V. Endotoxin: The uninvited guest. Biomaterials 26, 6811–6817. (2005)

30 Perrin-Cocon, L., Agaugué, S., Coutant, F., Masurel, A., Bezzine, S., Lambeau, G., André, P. & Lotteau, V. Secretory phospholipase A2 induces dendritic cell maturation. Eur J Immunol 34, 293–2302. (2004)

31 Okano, M., Nishizaki, K., Satoskar, A. R., Yoshino, T., Masuda, Y., & Harn, D. A. Jr. Involvement of carbohydrate on phospholipase A2, a bee-venom allergen, in in vivo antigen-specific IgE synthesis in mice. Allergy 54, 811–818 (1999).

32 Palm, N. W., Rosenstein, R. K., Yu, S., Schenten, D. D., Florsheim, E., & Medzhitov, R. Bee venom phospholipase A2 induces a primary type 2 response that is dependent on the receptor ST2 and confers protective immunity. Immunity 39, 976–985, doi:10.1016/j.immuni.2013.10.006 (2013).

33 Kitsiouli, E. I., Nakos, G. & Lekka, M. E. Differential determination of phospholipase A2 and PAF-acetylhydrolase in biological fluids using fluorescent substrates. J Lipid Res 40, 2346–2356. (1999)

34 Banks, W. A. Techniques for collecting, rearing, and handling imported fire ants. (1981).

35 Pitts, J. P., Hugh, M. C. J., & Ross, K. G. Cladistic analysis of the fire ants of the *Solenopsis saevissima* species-group (Hymenoptera: Formicidae). Zoologica Scripta, 34, 493–505. (2005)

36 Sobrinho, J. C., Kayano, A. M., Simões-Silva, R., Alfonso, J. J., Gomez, A. F., Gomez, M. C. V., Zanchi, F. B., Moura, L. A., Souza, V. R., Fuly, A. L., Oliveira, E., Silva, S. L., Almeida, J. R., Zuliani, J. P. & Soares, A. M. Antiplatelet aggregation activity of two novel acidic Asp49-phospholipases A2 from *Bothrops brazili* snake venom. Int J Biol Macrom, 107, 1014–1022. (2018).

